# AJILE12: Long-term naturalistic human intracranial neural recordings and pose

**DOI:** 10.1101/2021.07.26.453884

**Authors:** Steven M. Peterson, Satpreet H. Singh, Benjamin Dichter, Michael Scheid, Rajesh P. N. Rao, Bingni W. Brunton

## Abstract

Understanding the neural basis of human movement in naturalistic scenarios is critical for expanding neuroscience research beyond constrained laboratory paradigms. Here, we describe our *Annotated Joints in Long-term Electrocorticography for 12 human participants* (AJILE12) dataset, the largest human neurobehavioral dataset that is publicly available; the dataset was recorded opportunistically during passive clinical epilepsy monitoring. AJILE12 includes synchronized intracranial neural recordings and upper body pose trajectories across 55 semi-continuous days of naturalistic movements, along with relevant metadata, including thousands of wrist movement events and annotated behavioral states. Neural recordings are available at 500 Hz from at least 64 electrodes per participant, for a total of 1280 hours. Pose trajectories at 9 upper-body keypoints were estimated from 118 million video frames. To facilitate data exploration and reuse, we have shared AJILE12 on The DANDI Archive in the Neurodata Without Borders (NWB) data standard and developed a browser-based dashboard.

## Background & Summary

Natural human movements are complex and adaptable, involving highly coordinated sensorimotor processing in multiple cortical and subcortical areas^1–4^. However, many experiments focusing on the neural basis of human upper-limb movements often study constrained, repetitive motions such as center-out reaching within a controlled laboratory setup^5–9^. Such studies have greatly increased our knowledge about the neural correlates of movement, but it remains unclear how well these findings generalize to the natural movements that we often make in everyday situations^10,11^. Human upper-limb movement studies have incorporated self-cued and less restrictive movements^12–16^, but focusing on unstructured, naturalistic movements can enhance our knowledge of the neural basis of motor behaviors^17^, help us understand the role of neurobehavioral variability^18,19^, and aid in the development of robust brain-computer interfaces for real-world use^20–26^.

Here, we present synchronized intracranial neural recordings and upper body pose trajectories opportunistically obtained from 12 human participants while they performed unconstrained, naturalistic movements over 3–5 recording days each (55 days total). Intracranial neural activity, recorded via electrocorticography (ECoG), involves placing electrodes directly on the cortical surface, beneath the skull and dura, to provide high spatial and temporal resolution^27–29^. Pose trajectories were obtained from concurrent video recordings using computer vision to automate the often-tedious annotation procedure that has previously precluded the creation of similar datasets^30,31^. Along with these two core datastreams, we have added extensive metadata, including thousands of wrist movement initiation events previously used for neural decoding^32,33^, 10 quantitative event-related features describing the type of movement performed and any relevant context^18^, coarse labels describing the participant’s behavioral state based on visual inspection of videos^34^, and 14 different electrode-level features^18^. This dataset, which we call AJILE12 (*Annotated Joints in Long-term Electrocorticography for 12 human participants*), builds on our previous AJILE dataset^35^ and is depicted in Fig. 1.

**Figure 1.**
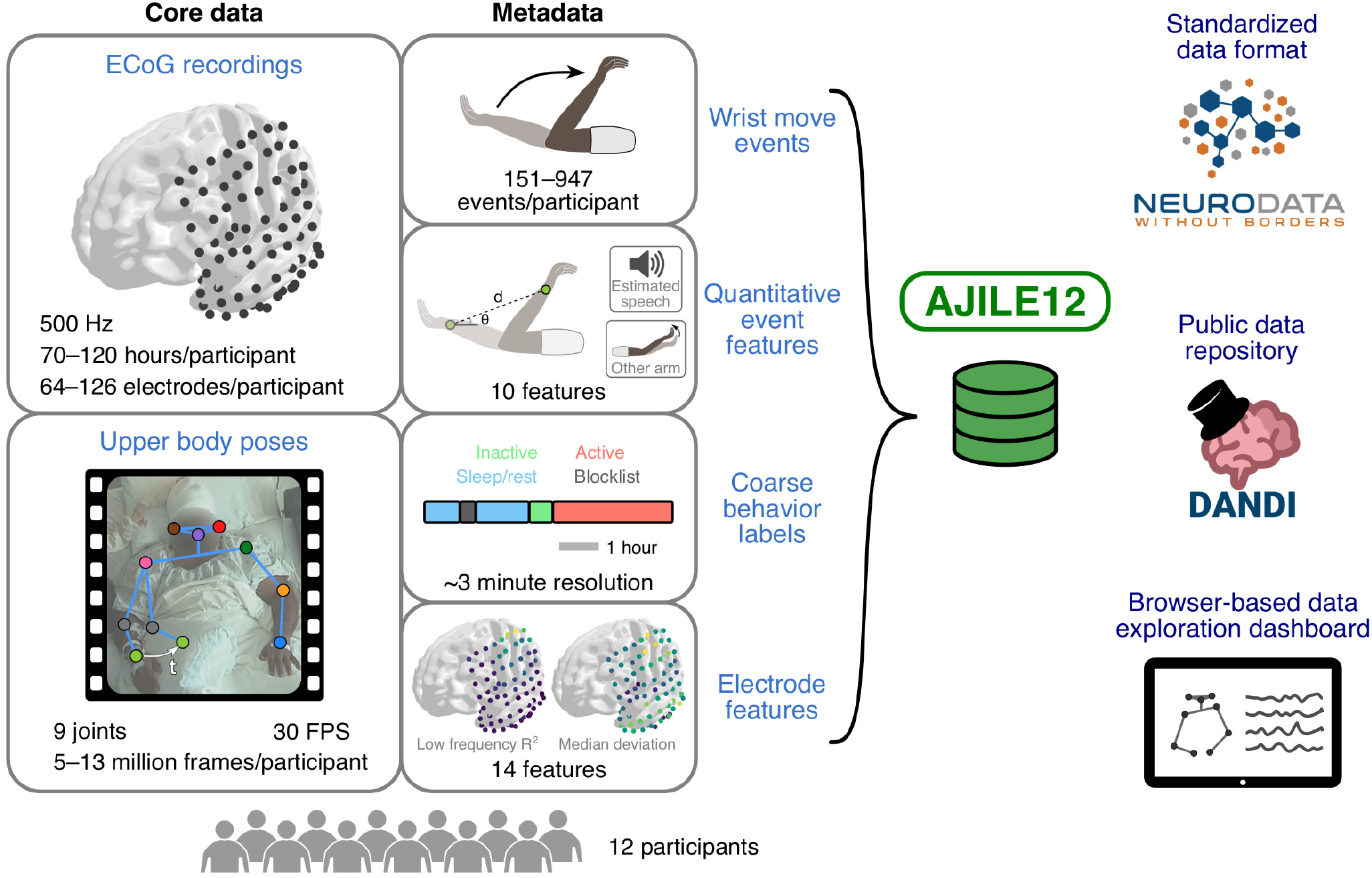
Schematic overview of our *Annotated Joints in Long-term Electrocorticography for 12 human participants* (AJILE12) dataset. AJILE12 includes ECoG recordings and upper body pose trajectories for 12 participants across 55 total recordings days, along with a variety of behavioral, movement event-related, and electrode-level metadata. All data is stored on The DANDI Archive in the NWB data standard, and we have created a custom browser-based dashboard in Jupyter Python to facilitate data exploration without locally downloading the data files.

AJILE12 has high reuse value for future analyses because it is large, comprehensive, well-validated, and shared in the NWB data standard. We have included 55 days of semi-continuous intracranial neural recordings along with thousands of verified wrist movement events, which both greatly exceed the size of typical ECoG datasets from controlled experiments^36^ as well as other long-term naturalistic ECoG datasets^34,35,37,38^. Such a wealth of data improves statistical power and enables large-scale exploration of more complex behaviors than previously possible, especially with modern machine learning techniques such as deep learning^32,39–42^. In addition, AJILE12 contains comprehensive metadata, including coarse behavior labels, quantitative event features, and localized electrode positions in group-level coordinates that enable cross-participant comparisons of neural activity. We have also pre-processed the neural data and visually validated all 6931 wrist movement events to ensure high-quality data, which have been already used in multiple studies^18,32,33^. In addition, we have released AJILE12 in the NWB data standard (Table 1)^43,44^ to adhere to the FAIR data principles of findability, accessibility, interoperability, and reusability^45^. Unified, open-source data formats such as NWB enable researchers to easily access the data and apply preexisting, reusable workflows instead of starting from scratch. Furthermore, we have developed an accessible and interactive browser-based dashboard that visualizes neural and pose activity, along with relevant metadata. This dashboard can access AJILE12 remotely to visualize the data without requiring local data file downloads, improving AJILE12’s accessibility.

**Table 1.** The main variables contained in each data file. Files are named sub-##_ses-#_behavior+ecephys.nwb, with ## indicating the participant and # denoting the day of recording.

| Data file variable | Description |
| --- | --- |
| \acquisition\ElectricalSeries | ECoG recordings |
| \processing\behavior\data_interfaces\Position | Upper body poses |
| \processing\behavior\data_interfaces\ReachEvents | Wrist move events |
| \intervals\reaches | Quantitative event features |
| \intervals\epochs | Coarse behavior labels |
| \electrodes | Electrode features |

## Methods

### Participants

We collected data from 12 human participants (8 males, 4 females; 29.4±7.6 years old [mean±SD]) during their clinical epilepsy monitoring at Harborview Medical Center (Seattle, USA). See Table 2 for individual participant details. Each participant had been implanted with electrocorticography (ECoG) electrodes placed based on clinical need. We selected these participants because they were generally active during their monitoring and had ECoG electrodes located near motor cortex. All participants provided written informed consent. Our protocol was approved by the University of Washington Institutional Review Board.

**Table 2.** Individual participant characteristics. Gender is denoted as male (M) or female (F), and implantation hemisphere is either left (L) or right (R). Surface electrodes denote grid and strip electrodes placed on the cortical surface, while depth electrodes reach deep cortical and subcortical areas. We identified bad/noisy electrodes based on high standard deviation or kurtosis values relative to that participant’s other electrodes. Good electrode counts are shown for each participant’s first available day of recording

| Participant | Gender | Age (years) | Recording days used | Hemisphere implanted | Surface electrodes: # good / total | Depth electrodes: # good / total |
| --- | --- | --- | --- | --- | --- | --- |
| P01 | M | 44 | 4 | L | 79 / 86 | 6 / 8 |
| P02 | M | 20 | 4 | R | 69 / 70 | 16 / 16 |
| P03 | M | 33 | 4 | L | 79 / 80 | 0 / 16 |
| P04 | F | 19 | 5 | R | 67 / 84 | 0 / 0 |
| P05 | F | 31 | 3 | R | 104 / 106 | 0 / 0 |
| P06 | M | 37 | 5 | L | 70 / 80 | 0 / 0 |
| P07 | M | 26 | 5 | R | 63 / 64 | 0 / 0 |
| P08 | F | 33 | 5 | R | 83 / 92 | 0 / 0 |
| P09 | M | 20 | 5 | L | 96 / 98 | 28 / 28 |
| P10 | M | 34 | 5 | L | 82 / 86 | 39 / 40 |
| P11 | F | 34 | 5 | L | 103 / 106 | 0 / 0 |
| P12 | M | 22 | 5 | L | 88 / 92 | 24 / 32 |

### Data collection

Semi-continuous ECoG and video were passively recorded from participants during 24-hour clinical monitoring for epileptic seizures. Recordings lasted 7.4±2.2 days (mean±SD) for each participant with sporadic breaks in monitoring (on average, 8.3±2.2 breaks per participant each lasting 1.9±2.4 hours). For all participants, we only included recordings during days 3–7 following the electrode implantation surgery to avoid potentially anomalous neural and behavioral activity immediately after the surgery. We excluded recording days with corrupted or missing data files, as noted in Table 2, and stripped all recording dates to de-identify participant data. These long-term, clinical recordings include various everyday activities, such as eating, sleeping, watching television, and talking while confined to a hospital bed. ECoG and video sampling rates were 1 kHz and 30 FPS (frames per second), respectively.

### ECoG data processing

We used custom MNE-Python scripts to process the raw ECoG data^46^. First, we removed DC drift by subtracting out the median voltage at each electrode. We then identified high-amplitude data discontinuities, based on abnormally high electrode-averaged absolute voltage (> 50 interquartile ranges [IQRs]), and set all data within 2 seconds of each discontinuity to 0.

With data discontinuities removed, we then band-pass filtered the data (1–200 Hz), notch filtered to minimize line noise at 60 Hz and its harmonics, downsampled to 500 Hz, and re-referenced to the common median for each grid, strip, or depth electrode group. For each recording day, noisy electrodes were identified based on abnormal standard deviation (> 5 IQRs) or kurtosis (> 10 IQRs) compared to the median value across electrodes. Using this procedure, we marked on average 7.3±5.6 ECoG electrodes as bad during each participant’s first available day of recording (Table 2).

Electrode positions were localized using the Fieldtrip toolbox in MATLAB. This process involved co-registering preoperative MRI and postoperative CT scans, manually selecting electrodes in 3D space, and warping electrode positions into MNI space (see Stolk et al.^47^ for further details).

### Markerless pose estimation

We performed markerless pose estimation on the raw video footage using separate DeepLabCut models for each participant^31^. First, we manually annotated the 2D positions of 9 upper-body keypoints (nose, ears, wrists, elbows, and shoulders) during 1000 random video frames for each participant (https://tinyurl.com/human-annotation-tool). Frames were randomly selected across all recording days, with preference towards frames during active, daytime periods. These manually annotated frames were used to train a separate DeepLabCut neural network model for each participant. We then applied the trained model to every video frame for that participant to generate estimated pose trajectories.

We synchronized ECoG data and pose trajectories using video timestamps and combined multiple recording sessions so that each file contained data from one entire 24-hour recording day that started and ended at midnight^48^.

### Wrist movement event identification

We used the estimated pose trajectories in order to identify unstructured movement initiation events of the wrist contralateral to the implanted hemisphere. To identify movement events, a first-order autoregressive hidden semi-Markov model was applied to the pose trajectory of the contralateral wrist. This model segmented the contralateral wrist trajectory into discrete move or rest states. Movement initiation events were identified as state transitions where 0.5 seconds of rest was followed by 0.5 seconds of wrist movement (see Singh et al.^33^ for further details).

Next, we selected the movement initiation events that most likely corresponded to actual reaching movements. We excluded arm movements during sleep, unrelated experiments, and private times based on coarse behavioral labels, which are described in the next section. In addition, we only retained movement events that (1) lasted between 0.5–4 seconds, (2) had DeepLabCut confidence scores > 0.4, indicating minimal marker occlusion, and (3) had parabolic wrist trajectories, as determined by a quadratic fit to the wrist’s radial movement (*R*^2^ > 0.6). We used this quadratic fit criteria to eliminate outliers with complex movement trajectories. For each recording day, we selected up to 200 movement events with the highest wrist speeds during movement onset. Finally, we visually inspected all selected movement events and removed those with occlusions or false positive movements (17.8%±9.9% of events [mean±SD]).

For each movement event, we also extracted multiple, quantitative behavioral and environmental features. To quantify movement trajectories, we defined a *reach* as the maximum radial displacement of the wrist during the identified movement event, as compared to wrist position at movement onset. Movement features include reach magnitude, reach duration, 2D vertical reach angle (90° for upward reaches, 90° for downward reaches), and radial speed during movement onset. We also include the recording day and time of day when each movement event occurred, as well as an estimate of speech presence during each movement using audio recordings.

In addition, we quantified the amount of bimanual movement for event based on ipsilateral wrist movement. These features include a binary classification of bimanual/unimanual based on temporal lag between wrist movement onsets, the ratio of ipsilateral to contralateral reach magnitude, and the amount of each contralateral move state that temporally overlapped with an ipsilateral move state. The binary feature was bimanual if at least 4 frames (0.13 seconds) of continuous ipsilateral wrist movement began either 1 second before contralateral wrist movement initiation or anytime during the contralateral wrist move state. Please see Peterson et al.^18^ for further methodological details.

### Coarse behavioral labels

To improve wrist movement event identification, we performed coarse annotation of the video recordings every ~3 minutes. These behavioral labels were either part of a blocklist to avoid during event detection or general activities/states that the participant was engaged in at the time. Identified activities include sleep/rest, inactive, and active behaviors, which were further subdivided into activities such as talking, watching TV, and using a computer or phone (Fig. 2). Blocklist labels include times where event detection would likely be inaccurate, such as camera movement and occlusion, as well as private times and unrelated research experiments. Some participants also have clinical procedure labels, indicating times when the clinical staff responded to abnormal participant behavior. We upsampled all labels to match the 30 Hz sampling rate of the pose data. Tables 3 and 4 show the duration of each label across participants for activity and blocklist labels, respectively.

**Figure 2.**
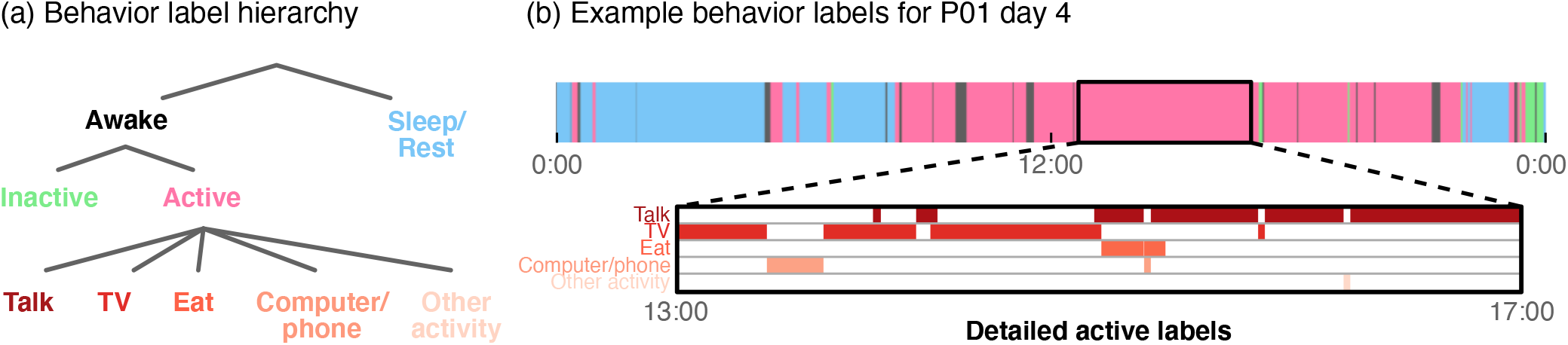
Coarse behavior labelling. (a) We annotated participant behavior in the video recordings using hierarchical labels to detail common awake and active behaviors. These annotations also include blocklist labels, which indicate times to potentially avoid during data exploration. (b) We show an example of the behavior labels for participant P01 during the entirety of recording day 4. Sleep/rest occurs in the morning and night times, as expected, with predominantly active periods during the day (8:00–20:00). Bottom row shows detailed active labels during a 4-hour active period that is dominated mostly by talk and TV behaviors. Note that these detailed active labels can overlap in time.

**Table 3.** Coarse activity label durations (in hours) for each participant. These labels describe various participant behaviors such as talking, eating, and watching television. Labels were generated by manual reviewing videos every ~3 minutes. While sleep is by far the most common, several activity labels appear over multiple hours for each participant. Note that multiple activity labels can co-occur (ex. eating while watching television). Therefore, the total duration of any activity label (last column) may be less than the sum of individual label durations for each participant.

| Participant | Sleep/rest | Inactive | Talk | TV | Computer/phone | Eat | Other activity | Total |
| --- | --- | --- | --- | --- | --- | --- | --- | --- |
| P01 | 29.2 | 1.9 | 15.6 | 22.0 | 7.8 | 2.6 | 0.8 | 75.7 |
| P02 | 53.2 | 2.1 | 9.2 | 7.7 | 2.9 | 2.1 | 4.8 | 79.8 |
| P03 | 11.9 | 0.3 | 21.8 | 1.9 | 19.8 | 3.5 | 0.9 | 51.1 |
| P04 | 34.0 | 3.1 | 28.3 | 11.0 | 8.2 | 2.0 | 1.2 | 78.9 |
| P05 | 35.4 | 2.8 | 8.9 | 12.1 | 5.8 | 1.0 | 1.2 | 62.5 |
| P06 | 37.8 | 0.1 | 2.8 | 5.7 | 2.6 | 0.3 | 0.2 | 45.3 |
| P07 | 46.8 | 0.3 | 5.0 | 0.3 | 1.9 | 0.4 | 0.2 | 53.6 |
| P08 | 47.8 | 0.8 | 6.8 | 5.3 | 1.3 | 0.6 | 2.2 | 61.8 |
| P09 | 87.3 | 8.1 | 3.8 | 0.0 | 0.0 | 0.6 | 1.9 | 100.6 |
| P10 | 67.5 | 2.1 | 5.8 | 0.1 | 5.6 | 0.0 | 1.8 | 81.4 |
| P11 | 36.0 | 1.4 | 6.6 | 0.0 | 0.1 | 0.0 | 0.8 | 44.9 |
| P12 | 32.4 | 0.3 | 1.5 | 0.1 | 0.5 | 0.0 | 0.6 | 35.4 |

**Table 4.** Coarse blocklist label durations (in hours) for each participant. These labels indicate times to avoid when extracting wrist movement events due to camera movements, unrelated experiments, and private times. Labels were generated by manual review of videos with ~3 minute resolution.

| Participant | Data break | Camera move/zoom | Camera occluded | Experiment | Private time | Tether/bandage | Hands under blanket | Clinical procedure | Total |
| --- | --- | --- | --- | --- | --- | --- | --- | --- | --- |
| P01 | 8.8 | 0.5 | 0.0 | 6.7 | 2.8 | 1.3 | 0.0 | 0.1 | 20.3 |
| P02 | 0.8 | 1.2 | 0.0 | 3.6 | 7.0 | 3.7 | 0.0 | 0.0 | 16.2 |
| P03 | 24.4 | 0.7 | 0.0 | 18.1 | 1.1 | 0.6 | 0.0 | 0.0 | 44.9 |
| P04 | 27.9 | 1.4 | 0.0 | 7.2 | 2.8 | 1.7 | 0.0 | 0.0 | 41.1 |
| P05 | 0.6 | 0.8 | 0.3 | 3.4 | 2.4 | 2.0 | 0.0 | 0.0 | 9.5 |
| P06 | 0.7 | 0.3 | 0.0 | 7.0 | 1.1 | 0.8 | 0.0 | 0.0 | 9.9 |
| P07 | 7.8 | 1.4 | 0.2 | 4.4 | 2.6 | 0.6 | 0.0 | 0.1 | 17.0 |
| P08 | 0.9 | 0.4 | 0.2 | 6.2 | 2.6 | 0.6 | 0.0 | 0.0 | 10.8 |
| P09 | 0.8 | 1.3 | 0.0 | 5.5 | 1.9 | 0.9 | 0.3 | 0.6 | 11.2 |
| P10 | 1.2 | 1.0 | 0.0 | 2.1 | 3.5 | 2.9 | 0.0 | 0.0 | 10.7 |
| P11 | 0.7 | 1.7 | 5.1 | 2.5 | 4.8 | 1.4 | 0.0 | 0.0 | 16.2 |
| P12 | 0.6 | 1.0 | 0.1 | 3.6 | 3.3 | 0.7 | 0.0 | 0.0 | 9.3 |

## Data Records

The data files are available on The DANDI Archive (https://identifiers.org/DANDI:000055)^48^, in the Neurodata Without Borders: Neurophysiology 2.0 (NWB:N) format^43,44^. All datastreams and metadata have been combined into a single file for each participant and day of recording, as indicated by the file name. For example, *sub-01_ses-3_behavior+ecephys.nwb* contains data from participant P01 on recording day 3. We used PyNWB 1.4.0 to load and interact with these data files. Table 1 shows the location of all main variables within each data file.

Each file contains continuous ECoG and pose data over a 24-hour period, with units of μV and pixels, respectively. ECoG data is located under \acquisition\ElectricalSeries as a *pynwb.ecephys.ElectricalSeries* variable. Pose data can be found under \processing\behavior\data_interfaces\Position as an *pynwb.behavior.Position* variable. Pose data is provided for the left/right ear (L_Ear, R_Ear), shoulder (L_Shoulder, R_Shoulder), elbow (L_Elbow, R_Elbow), and wrist (L_Wrist, R_Wrist), as well as the nose (Nose).

In addition to these core datastreams, each file contains relevant metadata. Contralateral wrist movement events are located in \processing\behavior\data_interfaces\ReachEvents as an *ndx_events.events.Events* variable. Quantitative neural and behavioral features for each event can be found in \intervals\reaches as a *pynwb.epoch.TimeIntervals* table with columns for each feature. Coarse behavioral labels are included in \intervals\epochs as a *pynwb.epoch.TimeIntervals* table. Each row contains the label along with the start and stop time in seconds.

We also include electrode-specific metadata in \electrodes as a *hdmf.common.table.DynamicTable*. Columns contain different metadata features, such as Montreal Neurological Institute (MNI) x, y, z coordinates and electrode group names. Electrode groups were named by clinicians based on their location in the brain. This table also contains the standard deviation, kurtosis, and median absolute deviation for each electrode computed over the entire recording file (excluding non-numeric values). Electrodes that we identified as noisy based on abnormal standard deviation and kurtosis are marked as False under the ‘good’ column. Table 2 shows the number of good electrodes that remain for each participant during the first available day of recording. We have also included the *R*^2^ scores obtained from regressing ECoG spectral power on the 10 quantitative event features for each participant’s wrist movement events^18^. Low-frequency power (used for low_freq_R2) indicates power between 8–32 Hz, while high-frequency power (used for high_freq_R2) denotes power between 76–100 Hz.

## Technical Validation

In this section, we assess the technical quality of AJILE12 by validating our two core datastreams: intracranial neural recordings and pose trajectories. In addition to this assessment, we have previously validated the quality and reliability of AJILE12 in multiple published studies^18,32,33^. We validated ECoG data quality by assessing spectral power projected into common brain regions^49^. This projection procedure enables multi-participant comparisons despite heterogeneous electrode coverage and reduces the dimensionality of the ECoG data from 64 or more electrodes (Fig. 3(a)) to a few brain regions of interest^18,32^. For this analysis, we focused on 4 sensorimotor and temporal regions in the left hemisphere defined using the AAL2 brain atlas^49,50^: precentral gyrus, postcentral gyrus, middle temporal gyrus, and inferior temporal gyrus. For participants with electrodes implanted primarily in the right hemisphere, we mirrored electrode positions into the left hemisphere. We divided the neural data into 30-minute windows and applied Welch’s method to compute the median spectral power over non-overlapping 30-second sub-windows^51^. We excluded 30-minute windows with non-numeric data values, likely due to data breaks. On average, we used 160.4±30.6 windows per participant (80.2±15.3 hours) across all recording days. Spectral power was interpolated to integer frequencies and projected into the 4 predefined brain regions (see Peterson et al.^18^ for further methodological details).

**Figure 3.**
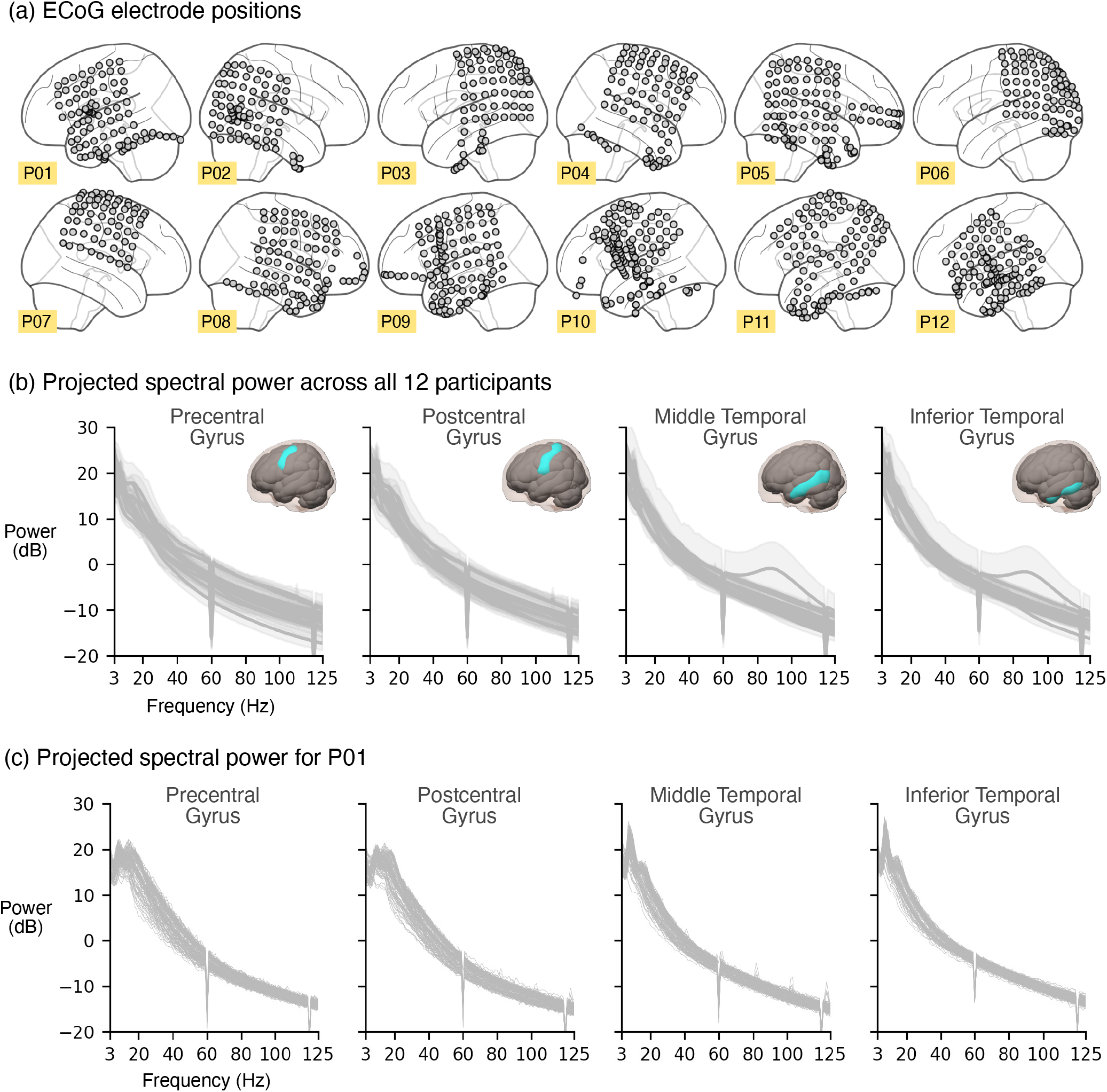
Validation of intracranial neural signal quality. (a) Electrocorticography (ECoG) electrode positions are shown in MNI coordinates for each participant. ECoG power spectra is shown for (b) all 12 participants (shading denotes standard deviation) and (c) participant P01 over all available half-hour time windows. We projected spectral power into sensorimotor and temporal brain regions, excluding time windows with non-numeric values that likely indicated a data break. Lines for participant P01 denote power in each window (*n* = 130 total, or 65 hours). The power spectra shape (exponential decrease for increasing frequencies) and consistency over time demonstrate the cleanliness and stability of our neural recordings across multiple recording days.

Fig. 3(b) shows the average spectral power across time windows, separated by participant. In general, power spectra remain quite consistent across participants with tight standard deviations across time windows, indicating that much of the ECoG data is good to use^52,53^. We also plotted the power spectra of each individual window for participant P01, as shown in Fig. 3(c). Again, the variation among time windows appears small, and we see clear differences in spectral power between sensorimotor (pre/postcentral gyri) and temporal areas, as expected. Additionally, we retained 92.3%±6.3% ECoG electrodes per participant (Table 2), further demonstrating the quality of our neural data^54,55^.

We validated pose trajectories by comparing each pose estimation model’s output to our manual annotations of each participant’s pose (Table 5). While manual annotations are susceptible to human error^56^, they are often used to evaluate markerless pose estimation performance when marker-based motion capture is not possible^30,57^. We used root-mean-square (RMS) error averaged across all keypoints to evaluate model performance for the 950 frames used to train the model as well as 50 annotated frames that were withheld from training. RMS errors for the holdout set (5.71±1.90 pixels) are notably larger than the train set errors (1.52±0.12 pixels), as expected, but are still within an acceptable tolerance given that 3 pixels are approximately equal to just 1 cm^33^.

**Table 5.** Pose estimation model errors. Root-mean-square reconstruction errors of our automated markerless pose models are shown for each participant’s train (950 frames) and holdout (50 frames) sets. We used manual annotations as ground truth when computing the error, which was averaged across all 9 upper-body keypoints. For reference, 3 pixels are approximately equal to 1 cm.

| Participant | Train set<br>error (pixels) | Holdout set<br>error (pixels) |
| --- | --- | --- |
| P01 | 1.45 | 4.27 |
| P02 | 1.44 | 3.58 |
| P03 | 1.58 | 6.95 |
| P04 | 1.63 | 6.02 |
| P05 | 1.43 | 3.42 |
| P06 | 1.43 | 6.63 |
| P07 | 1.51 | 5.45 |
| P08 | 1.84 | 10.35 |
| P09 | 1.40 | 4.05 |
| P10 | 1.48 | 7.59 |
| P11 | 1.51 | 5.45 |
| P12 | 1.52 | 4.73 |

## Usage Notes

We have developed a Jupyter Python dashboard that can be run online to facilitate data exploration without locally downloading the data files (https://github.com/BruntonUWBio/ajile12-nwb-data). Our dashboard includes visualizations of electrode locations, along with ECoG and wrist pose traces for a user-selected time window (Fig. 4). Users can also visualize the average contralateral wrist trajectory during identified movement events for each file. The dashboard streams from The DANDI Archive only the data needed for visualization, enabling efficient renderings of time segments from the large, 24-hour data files. Our code repository also includes all scripts necessary to create Figs. 2–3 and Tables 2–4. In addition, we have previously used AJILE12 to decode and analyze the neurobehavioral variability of naturalistic wrist movements and have publicly released multiple workflows that can be modified for use on this dataset^18,32,33^.

**Figure 4.**
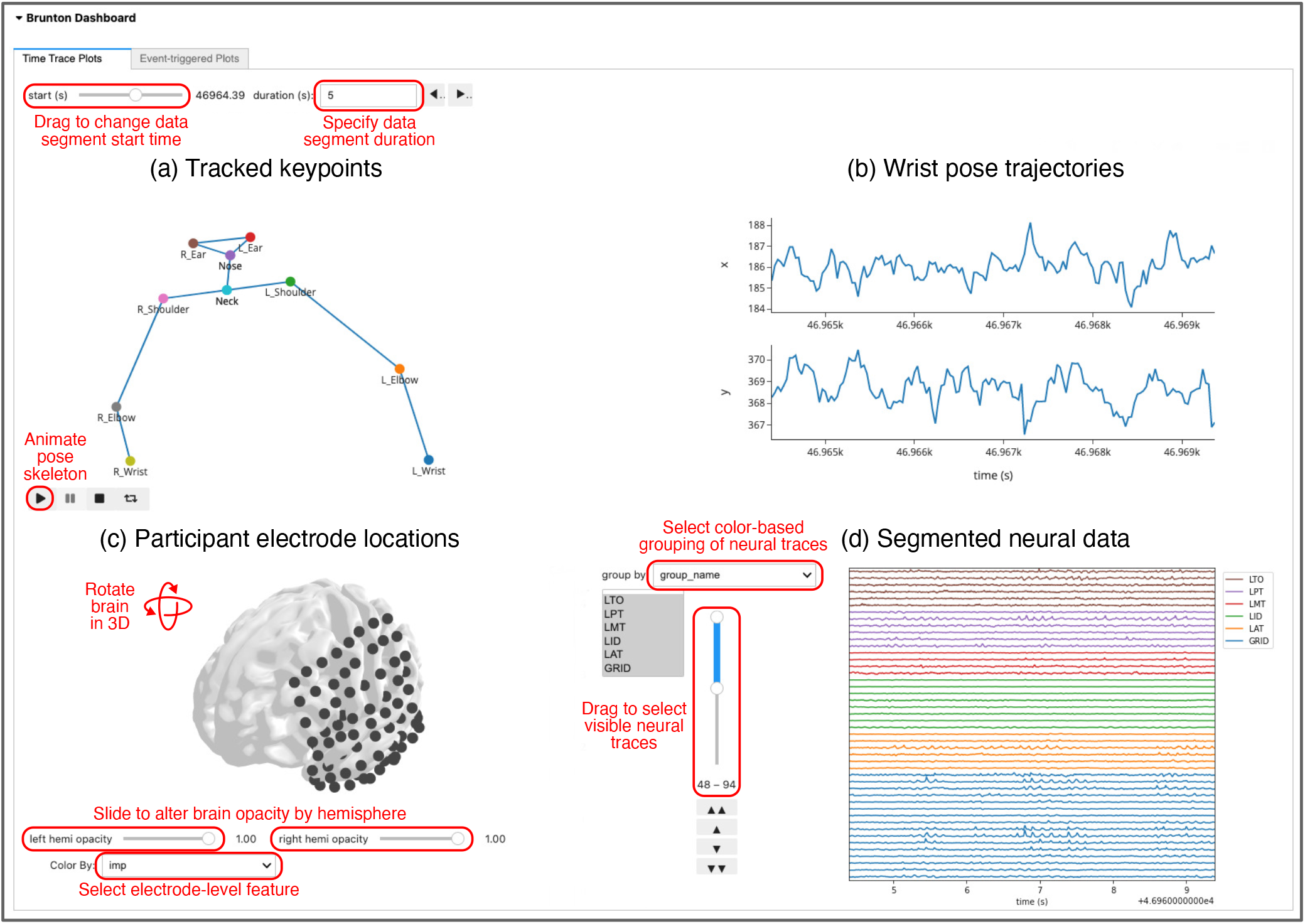
Browser-based Jupyter Python dashboard for dataset exploration. We designed a browser-based dashboard, available at https://github.com/BruntonUWBio/ajile12-nwb-data, to facilitate exploration of AJILE12 without needing to download any data files locally. (a) Participant keypoint positions are displayed for the first sample of a user-defined time window, with the option to animate keypoint positions across the entire window. We included a virtual neck marker for this visualization at the midpoint between the left and right shoulders. (b) Time-series traces of horizontal (x) and vertical (y) wrist positions are displayed over the same selected time window. (c) Electrode coverage is shown in MNI coordinates on a standardized brain model. This visualization is interactive, allowing three-dimensional rotations, alterations of hemisphere opacity to inspect depth electrodes, and the ability to visualize various electrode-level metadata such as electrode groups and identified bad electrodes. (d) Raw ECoG signals are visualized over the same user-selected time window, color-coded by electrode group.

## Code availability

Code to run our Jupyter Python dashboard and recreate all results in this paper can be found at https://github.com/BruntonUWBio/ajile12-nwb-data. We used Python 3.8.5 and PyNWB 1.4.0. A requirements file listing the Python packages and versions necessary to run the code is provided in our code repository. Our code is publicly available without restriction other than attribution.

## Acknowledgements

We thank Nancy Wang for contributing to the data collection, John So for generating the coarse behavior annotations, and the clinical staff at the Harborview Hospital Neurosurgery department for their assistance collecting and analyzing the data, especially Leigh Weber, Jeffrey G. Ojemann, and Andrew Ko. This research was supported by funding from the National Science Foundation (1630178 and EEC-1028725), the Defense Advanced Research Projects Agency (FA8750-18-2-0259), the Sloan Foundation, the Washington Research Foundation, and the Weill Neurohub.

## Author contributions statement

R.P.N.R and B.W.B conceived the study; S.M.P. and S.H.S. performed the data analysis; S.M.P., S.H.S., R.P.N.R, and B.W.B. interpreted the results; S.M.P., B.D., and M.S. created the public dataset and corresponding analysis dashboard; S.M.P. and B.W.B. wrote the paper; all authors reviewed and approved the final draft of the paper.

